# A benzoxazolylurea inhibits VraS and enhances antimicrobials against vancomycin intermediate-resistant *Staphylococcus aureus*

**DOI:** 10.1101/2023.08.31.555806

**Authors:** Emerson P. Heckler, Liaqat Ali, Shrijan Bhattarai, Brittnee Cagle-White, Nicklaus Smith, Robert A. Coover, May H. Abdel Aziz, Aurijit Sarkar

## Abstract

Vancomycin intermediate-resistant Staphylococcus aureus (VISA) is a pathogen of concern. VraS, a histidine kinase, facilitates the VISA phenotype. Here, we reveal a benzoxazolylurea that directly inhibits VraS and enhances vancomycin to below the clinical breakpoint against an archetypal VISA strain. To the best of our knowledge, this is the first direct VraS inhibitor ever reported that shows significant enhancement of vancomycin against VISA.

---

The rise of antibiotic resistance not only affects our ability to prevent and treat infections, but also threatens the viability of several contemporary medical procedures, such as surgeries, organ transplants and anticancer chemotherapy (1, 2). Finding novel antimicrobials has been difficult due to a complex set of circumstances, including scientific and financial hurdles (3-6). A potential solution to the problem is to retain the clinical efficacy of our current arsenal of antimicrobials. Amoxiclav is a clinically used example of this approach against penicillinase-expressing, methicillin-susceptible *Staphylococcus aureus* (MSSA). No such options exist against methicillin-resistant (MRSA) and vancomycin intermediate-resistant (VISA) *S. aureus*.

Beta-lactams and glycopeptides are preferred treatments for various staphylococcal infections. Oxacillin is a first-line treatment for MSSA infections, but vancomycin is the drug-of-choice against MRSA infections (7) because oxacillin is ineffective. VISA strains cannot be treated by vancomycin or oxacillin (8). Restoring the clinical efficacy of either oxacillin or vancomycin (or both) is desirable.

Vancomycin shows a minimum inhibitory concentration (MIC) of ∼4 to 8 μg/mL against VISA, whereas the clinical breakpoint is 2 μg/mL (9). A thicker cell wall in VISA enables resistance (10-12). VraS, of VraSRT, encoded by the *vra* operon, is a critical contributor to the VISA phenomenon (13-22). Enhanced VraSR signaling in a susceptible *S. aureus* strain increases the MIC of vancomycin to intermediate-resistant levels (19). Cell wall stress activates VraS signaling (23), leading to auto-upregulation and activation of the cell wall stress stimulon (24) in an attempt to repair the protective structure. VraS thus gains prominence as a potential drug target associated with beta-lactam and glycopeptide resistance (15, 25-27). Mutational inactivation of *vra* signaling reduces MIC of oxacillin to below the accepted breakpoint of 4 μg/mL in MRSA (28). VraS is a validated target because its inhibitors are powerful enhancers of vancomycin and oxacillin against *S. aureus* (27, 29, 30).

We have identified a powerful VraS inhibitor, chemical ***1*** (**Fig 1**) along with a preliminary structure-function relationship study. An NMR spectrum (**Figure S1**) ascertained the chemical’s identity.

**Figure 1:**
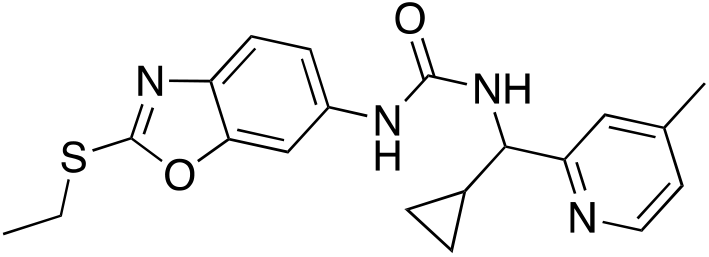
Chemical ***1***, IUPAC name: 1-(cyclopropyl(4-methylpyridin-2-yl)methyl)-3-(2-(ethylthio)benzo[d]oxazol-6-yl)urea, a urea, enhances vancomycin and oxacillin against VISA.

We ran a screen for vancomycin enhancers against Mu50 (8), an archetypal VISA strain (see (31) and **Supplementary Methods** for details). Mu50 was intermediate-resistant (vancomycin MIC 4 μg/mL) as expected (9). A USA300 MRSA, ATCC BAA-1717, was susceptible at 2 μg/mL (32). Chemical ***1*** enhanced vancomycin activity by 16-fold against Mu50 (**Table 1**). The MIC of vancomycin dropped to 0.25 μg/mL when we added ***1*** at a final concentration of 50 μM. Oxacillin MIC changed from 512 μg/mL to 8 μg/mL. Penicillin G and chloramphenicol showed minimal or no change in MIC. Enhancement of vancomycin was minimal against the MRSA strain, comparable with previous reports (29), but oxacillin and penicillin G were not enhanced. Chloramphenicol MIC reduction was borderline as well. Interestingly, two structurally similar chemicals (**Fig S2**) did not show similar activity.

**Table 1:**
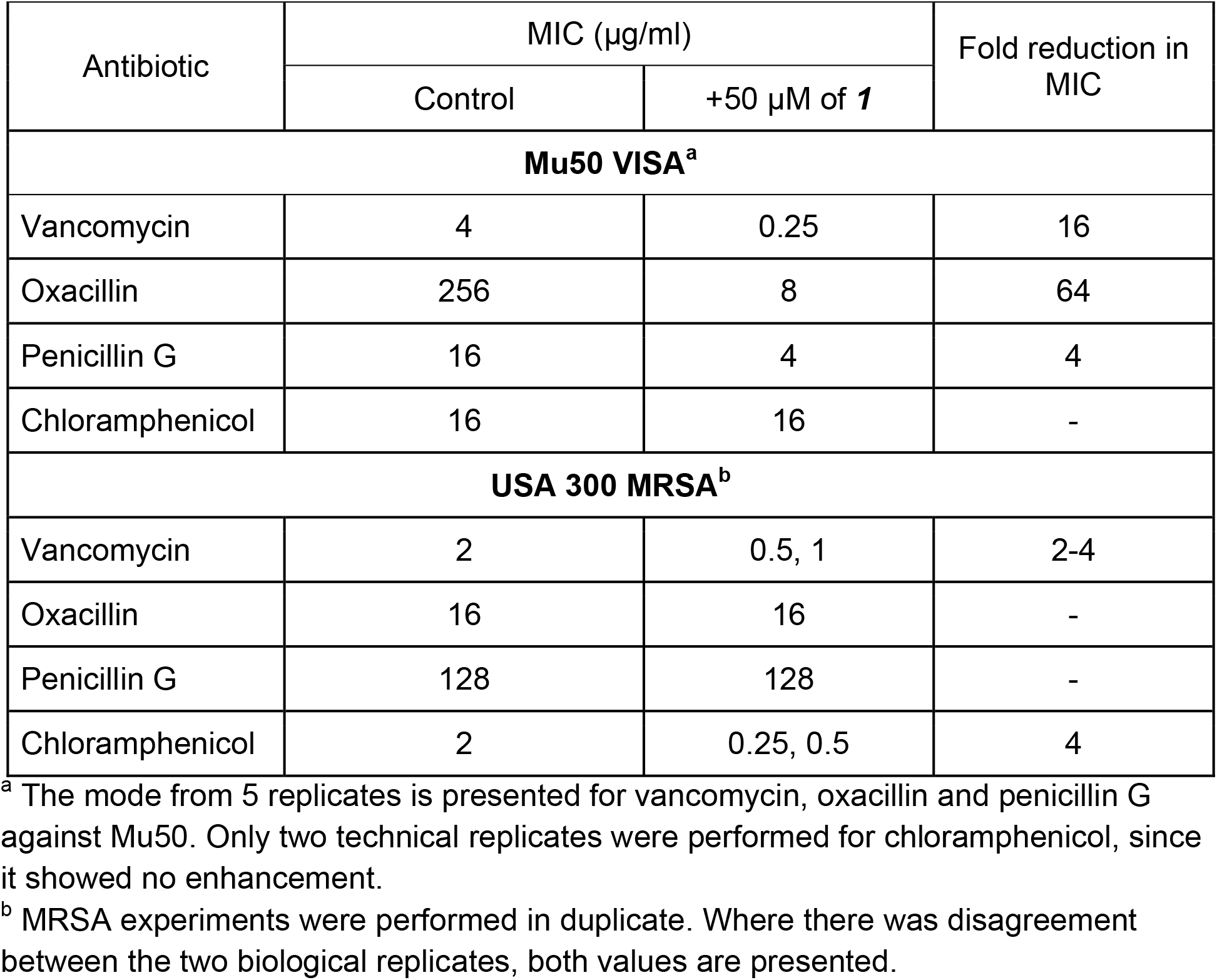
MIC of antibiotics vs. USA 300 and Mu50 with and without ***1***.

**Fig 2** shows Fractional Inhibitory Concentration Index (FICI) values. The MIC of ***1*** alone was >400 μM, so the chemical shows synergy with vancomycin (FICI ≤ 0.12), oxacillin (FICI ≤ 0.31) and penicillin G (FICI ≤ 0.31) against Mu50 VISA.

**Figure 2:**
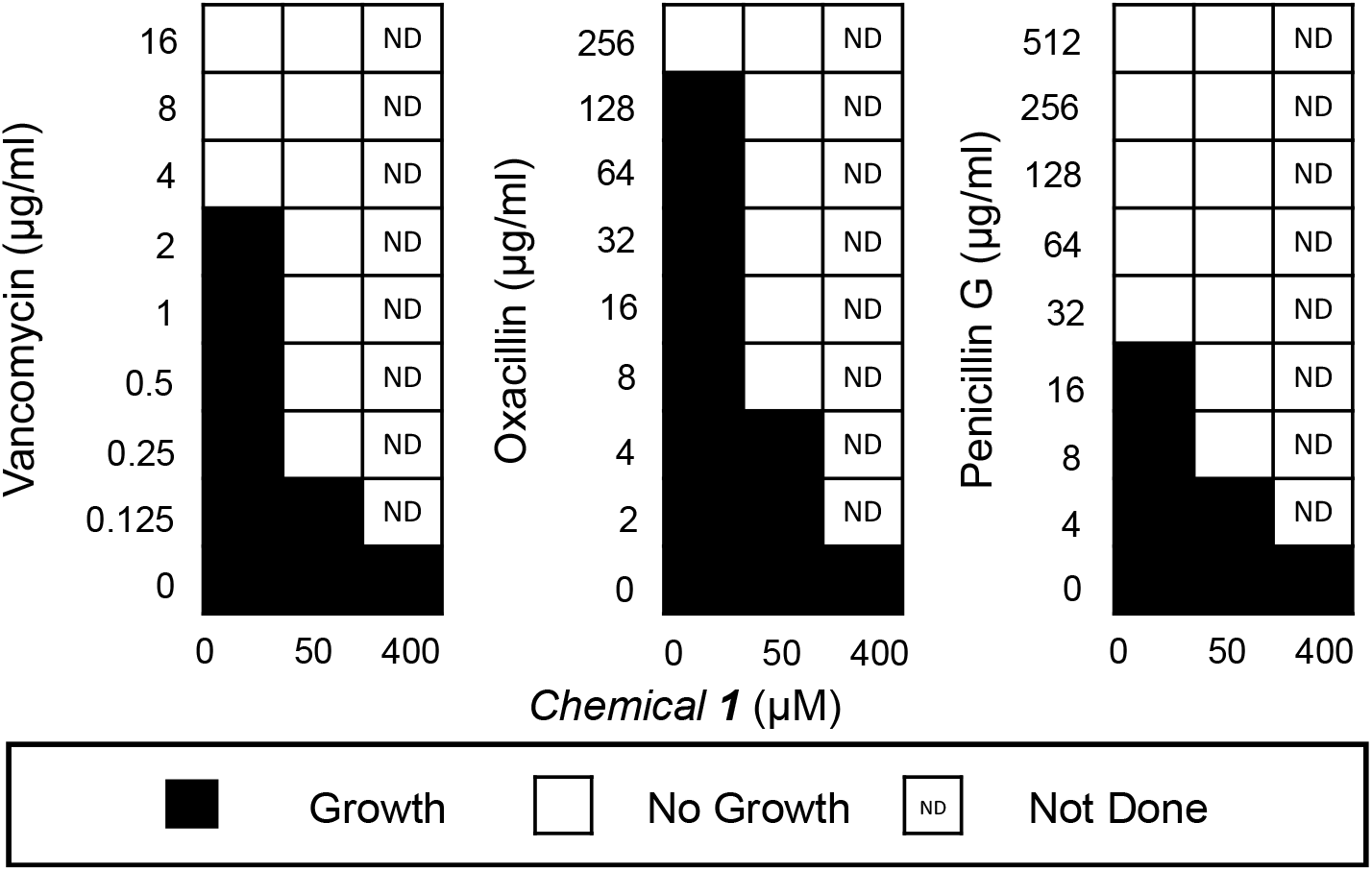
Checkerboard assays demonstrate that ***1*** synergizes with vancomycin, oxacillin and penicillin G against VISA.

The *vra* operon is central to resistance against cell wall-acting antimicrobial (13-15, 21-23, 25, 27, 28). Antibiotic stress auto-upregulates *vraSR* and *vraS* inactivation blocks this auto-upregulation (28). Chemical inhibition of VraS also reduces *vraSR* transcript levels (27, 30). We monitored *vraSR* transcript levels in Mu50 using RT-qPCR in presence of ***1*** (**Fig 3**) as previously described (30). See **Supplementary Methods** for details. *vraSR* transcript levels upon exposure to ***1*** alone were indistinguishable from untreated Mu50. When Mu50 was exposed to vancomycin or oxacillin, *vraSR* transcript levels increased ∼6-fold, but failed to increase as much when ***1*** was present. Thus, *vra* auto-upregulation is inhibited by ***1***.

**Figure 3:**
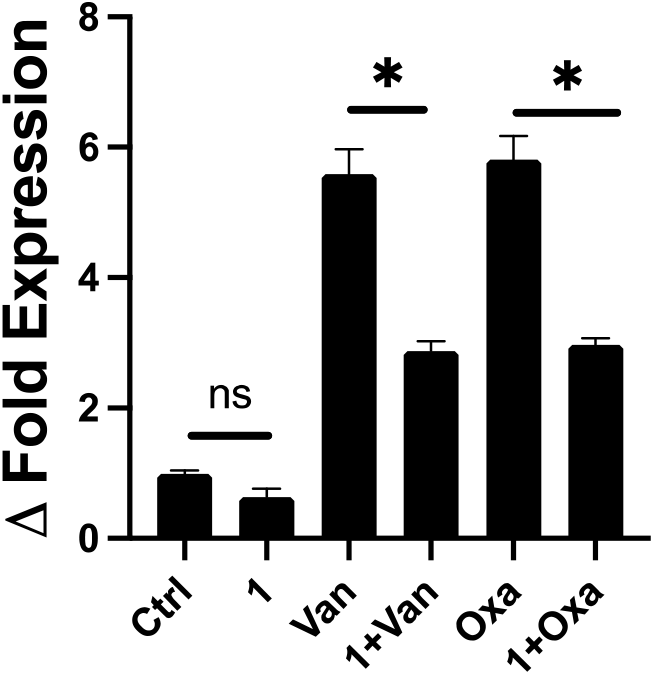
Monitoring *vraSR* auto-upregulation under antibiotic stress (Ctrl: no treatment; Van: vancomycin; Oxa: oxacillin), with and without ***1***, using RT-qPCR.

We tested direct inhibition of purified GST-VraS with ***1***. GST-VraS was expressed and purified with >90% purity and protein identity confirmed by Western blots (WB) using a tag-specific antibody as previously reported (14). The rate of the ATPase activity of GST-VraS (1 μM) was assessed using a kinase-coupled assay in the presence of increasing concentrations of ***1*** (0.1 – 300 μM) and analyzed to determine the IC50 as previously described (30). Briefly, the conversion of ATP to ADP is coupled to the oxidation of NADH in the pyruvate kinase/lactate dehydrogenase system. The reaction of the ATPase corresponds to the decrease in the fluorescent signal of NADH (λexcitation 340 nm, λemission 455 nm). ***1*** was found to inhibit the ATPase activity of GST-VraS with an IC50 of 0.94 ± 0.29 μM. The enzyme activity was inhibited by a maximum of 35%, indicating residual activity at the highest inhibitor concentration (**Fig 4**). It is therefore unequivocally demonstrated that VraS is a direct target of ***1***.

**Figure 4:**
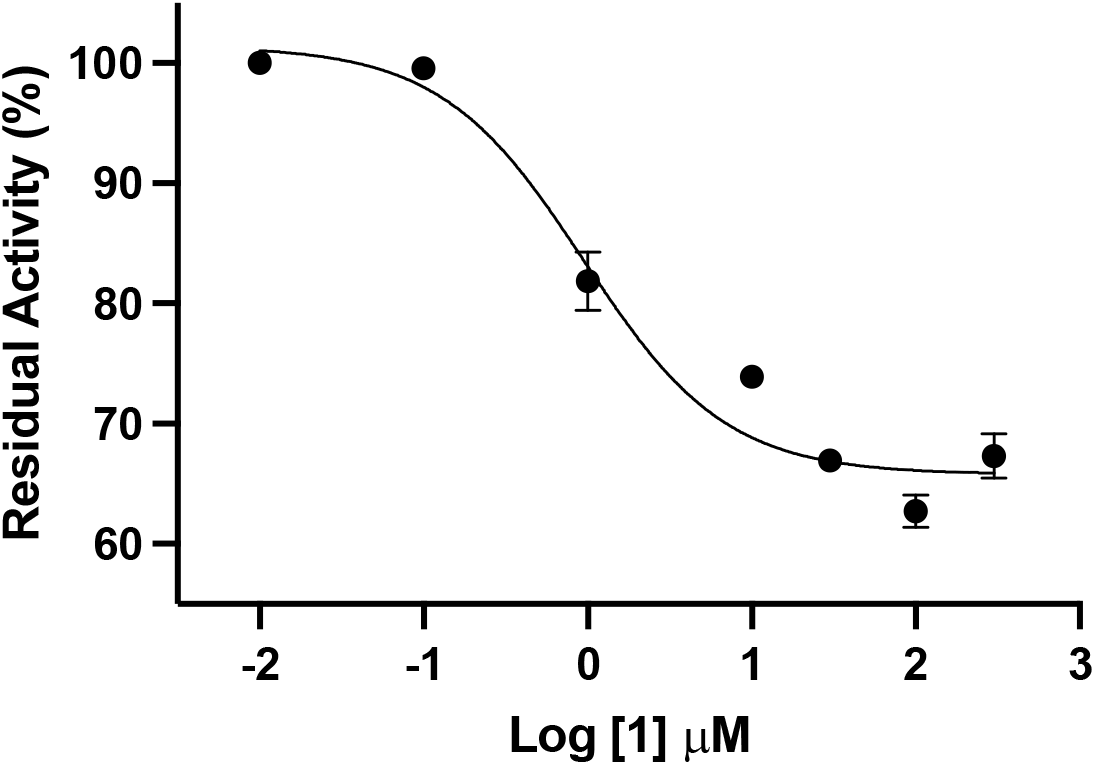
Direct inhibition of VraS by ***1***.

The crystal structure of the VraS catalytic domain with ADP bound in its active site is available (RCSB Protein Data Bank code *4gt8*). We developed a docking protocol based on it (see **Supplementary Methods** for details). VraS structure was prepared using UCSF Chimera (33). ADP was extracted to vacate the binding site. The structures of ADP, adenosine and ***1*** were prepared. Adenosine was chosen as a more hydrophobic ADP analog; Docking parameters are likely relevant for drug-like chemicals if the adenine ring in adenosine and ADP dock successfully. Both stereoisomers of ***1*** were modeled since our experimental procedures did not use a specific stereoisomer.

All three were docked (**Fig 5**) in 3 independent runs using the PLANTS algorithm (34, 35). ADP and adenosine docked reproducibly, but neither stereoisomer of chemical ***1*** did (not shown), suggesting ***1*** is likely an allosteric VraS inhibitor, like others (30).

**Figure 5:**
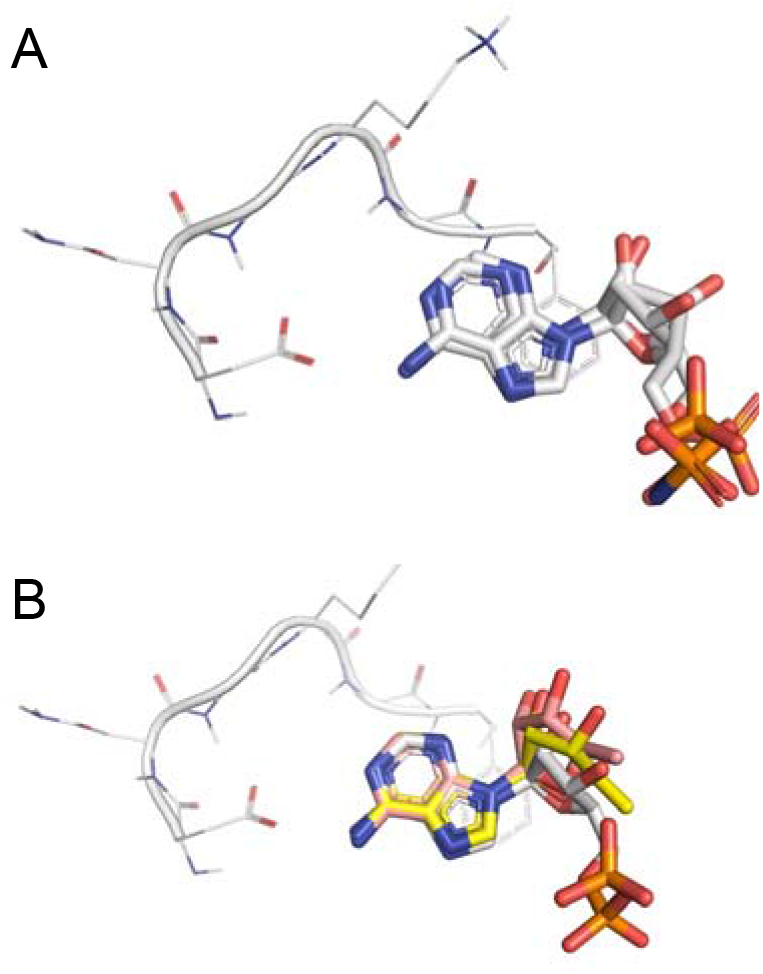
A validated docking protocol to find ATP-competitive inhibitors of VraS. While **(A)** ADP and **(B)** adenosine reproducibly docked into the VraS structure (RCSB PDB code 4gt8, chain A) in triplicate runs. Adenosine is shown overlayed on the co-crystallized ligand, ADP.

Previous studies on VraS inhibitors focused on oxacillin enhancement against *S. aureus* (27, 29). Our study reports enhancement of both, vancomycin and oxacillin. Previous urea inhibitors of VraS (27) were structurally distinct (phenylurea is the maximum common substructure), although of comparable potency in oxacillin enhancement and VraS binding affinity. The activity of ***1*** is yet to be optimized, but is already within a few fold of the best known VraS inhibitors (27, 29). VISA appears susceptible to vancomycin and near-susceptible to oxacillin, when exposed to ***1***. Therefore, ***1*** is a very interesting scaffold in our hunt for enhancers of vancomycin and oxacillin against VISA.

## SUPPLEMENTAL INFORMATION

The supporting information is available free of charge at the online version of this manuscript. This includes characterization details for chemicals, all replicates of antimicrobial susceptibility assessment, and methodological details to help with replication of this work.

## AUTHOR INFORMATION

### Author contributions

Conceived the work: AS, MHA

Performed the work: EPH, BC-W, LA, SB, NS

Analyzed the data: all authors

Wrote/edited the manuscript: all authors

All authors have reviewed the manuscript

## ACKNOWLEDGEMENTS

This work was supported by American Association of Colleges of Pharmacy New Investigator Award and American Heart Association award no. 959516 to AS. The authors would also like to acknowledge salary support by The University of Texas at Tyler to LA.

## NOTES

The authors declare no competing financial interests.

## REFERENCES

1. Sarkar A, Garneau-Tsodikova S. 2019. Resisting resistance: gearing up for war. MedChemCommun 10:1512–1516.

2. Brown ED, Wright GD. 2016. Antibacterial drug discovery in the resistance era. Nature 529:336–43.

3. O’Shea R, Moser HE. 2008. Physicochemical properties of antibacterial compounds: implications for drug discovery. Journal of medicinal chemistry 51:2871–8.

4. Gwynn MN, Portnoy A, Rittenhouse SF, Payne DJ. 2010. Challenges of antibacterial discovery revisited. Ann N Y Acad Sci 1213:5–19.

5. Power E. 2006. Impact of antibiotic restrictions: the pharmaceutical perspective. Clin Microbiol Infect 12 Suppl 5:25–34.

6. Payne DJ, Gwynn MN, Holmes DJ, Pompliano DL. 2007. Drugs for bad bugs: confronting the challenges of antibacterial discovery. Nat Rev Drug Discov 6:29–40.

7. Tong SY, Davis JS, Eichenberger E, Holland TL, Fowler VG, Jr. 2015. Staphylococcus aureus infections: epidemiology, pathophysiology, clinical manifestations, and management. Clin Microbiol Rev 28:603–61.

8. Hiramatsu K, Hanaki H, Ino T, Yabuta K, Oguri T, Tenover FC. 1997. Methicillin-resistant Staphylococcus aureus clinical strain with reduced vancomycin susceptibility. J Antimicrob Chemother 40:135–6.

9. Howden BP, Davies JK, Johnson PD, Stinear TP, Grayson ML. 2010. Reduced vancomycin susceptibility in Staphylococcus aureus, including vancomycin-intermediate and heterogeneous vancomycin-intermediate strains: resistance mechanisms, laboratory detection, and clinical implications. Clin Microbiol Rev 23:99–139.

10. Hanaki H, Kuwahara-Arai K, Boyle-Vavra S, Daum RS, Labischinski H, Hiramatsu K. 1998. Activated cell-wall synthesis is associated with vancomycin resistance in methicillin-resistant Staphylococcus aureus clinical strains Mu3 and Mu50. Journal of Antimicrobial Chemotherapy 42:199–209.

11. Cui L, Murakami H, Kuwahara-Arai K, Hanaki H, Hiramatsu K. 2000. Contribution of a thickened cell wall and its glutamine nonamidated component to the vancomycin resistance expressed by Staphylococcus aureus Mu50. Antimicrob Agents Chemother 44:2276–85.

12. Cui L, Ma X, Sato K, Okuma K, Tenover FC, Mamizuka EM, Gemmell CG, Kim MN, Ploy MC, El-Solh N, Ferraz V, Hiramatsu K. 2003. Cell wall thickening is a common feature of vancomycin resistance in Staphylococcus aureus Journal of Clinical Microbiology 41:5–14.

13. Boyle-Vavra S, Yin S, Jo DS, Montgomery CP, Daum RS. 2013. VraT/YvqF is required for methicillin resistance and activation of the VraSR regulon in Staphylococcus aureus. Antimicrob Agents Chemother 57:83–95.

14. Belcheva A, Golemi-Kotra D. 2008. A close-up view of the VraSR two-component system. A mediator of Staphylococcus aureus response to cell wall damage. J Biol Chem 283:12354–64.

15. Gardete S, Wu SW, Gill S, Tomasz A. 2006. Role of VraSR in antibiotic resistance and antibiotic-induced stress response in Staphylococcus aureus. Antimicrob Agents Chemother 50:3424–34.

16. Sieradzki K, Tomasz A. 2003. Alterations of Cell Wall Structure and Metabolism Accompany Reduced Susceptibility to Vancomycin in an Isogenic Series of Clinical Isolates of Staphylococcus aureus. Journal of Bacteriology 185:7103–7110.

17. Qureshi NK, Yin S, Boyle-Vavra S. 2014. The role of the Staphylococcal VraTSR regulatory system on vancomycin resistance and vanA operon expression in vancomycin-resistant Staphylococcus aureus. PLoS One 9:e85873.

18. Matsuo M, Cui L, Kim J, Hiramatsu K. 2013. Comprehensive identification of mutations responsible for heterogeneous vancomycin-intermediate Staphylococcus aureus (hVISA)-to-VISA conversion in laboratory-generated VISA strains derived from hVISA clinical strain Mu3. Antimicrob Agents Chemother 57:5843–53.

19. Kuroda M, Kuwahara-Arai K, Hiramatsu K. 2000. Identification of the up- and down-regulated genes in vancomycin-resistant Staphylococcus aureus strains Mu3 and Mu50 by cDNA differential hybridization method. Biochem Biophys Res Commun 269:485–90.

20. Kato Y, Suzuki T, Ida T, Maebashi K, Sakurai M, Shiotani J, Hayashi I. 2008. Microbiological and clinical study of methicillin-resistant Staphylococcus aureus (MRSA) carrying VraS mutation: changes in susceptibility to glycopeptides and clinical significance. Int J Antimicrob Agents 31:64–70.

21. Galbusera E, Renzoni A, Andrey DO, Monod A, Barras C, Tortora P, Polissi A, Kelley WL. 2011. Site-specific mutation of Staphylococcus aureus VraS reveals a crucial role for the VraR-VraS sensor in the emergence of glycopeptide resistance. Antimicrob Agents Chemother 55:1008–20.

22. Cui L, Neoh HM, Shoji M, Hiramatsu K. 2009. Contribution of vraSR and graSR point mutations to vancomycin resistance in vancomycin-intermediate Staphylococcus aureus. Antimicrob Agents Chemother 53:1231–4.

23. McCallum N, Meier PS, Heusser R, Berger-Bachi B. 2011. Mutational analyses of open reading frames within the vraSR operon and their roles in the cell wall stress response of Staphylococcus aureus. Antimicrob Agents Chemother 55:1391–402.

24. Utaida S, Dunman PM, Macapagal D, Murphy E, Projan SJ, Singh VK, Jayaswal RK, Wilkinson BJ. 2003. Genome-wide transcriptional profiling of the response of Staphylococcus aureus to cell-wall-active antibiotics reveals a cell-wall-stress stimulon. Microbiology 149:2719–2732.

25. Kuroda M, Kuroda H, Oshima T, Takeuchi F, Mori H, Hiramatsu K. 2004. Two-component system VraSR positively modulates the regulation of cell-wall biosynthesis pathway in Staphylococcus aureus. Molecular Microbiology 49:807–821.

26. Yin S, Daum RS, Boyle-Vavra S. 2006. VraSR two-component regulatory system and its role in induction of pbp2 and vraSR expression by cell wall antimicrobials in Staphylococcus aureus. Antimicrobial Agents Chemotherapy 50:336–43.

27. Lee H, Boyle-Vavra S, Ren J, Jarusiewicz JA, Sharma LK, Hoagland DT, Yin S, Zhu T, Hevener KE, Ojeda I, Lee RE, Daum RS, Johnson ME. 2019. Identification of Small Molecules Exhibiting Oxacillin Synergy through a Novel Assay for Inhibition of vraTSR Expression in Methicillin-Resistant Staphylococcus aureus. Antimicrob Agents Chemother 63.

28. Boyle-Vavra S, Yin S, Daum RS. 2006. The VraS/VraR two-component regulatory system required for oxacillin resistance in community-acquired methicillin-resistant Staphylococcus aureus. FEMS Microbiol Lett 262:163–71.

29. Harris TL, Worthington RJ, Melander C. 2012. Potent small-molecule suppression of oxacillin resistance in methicillin-resistant Staphylococcus aureus. Angew Chem Int Ed Engl 51:11254–7.

30. Bhattarai S, Marsh L, Knight K, Ali L, Gomez A, Sunderhaus A, Abdel Aziz MH. 2023. NH125 Sensitizes Staphylococcus aureus to Cell Wall-Targeting Antibiotics through the Inhibition of the VraS Sensor Histidine Kinase. Microbiol Spectr 11:e0486122.

31. Thomas PM, Deming MA, Sarkar A. 2022. β-Lactamase Suppression as a Strategy to Target Methicillin-Resistant Staphylococcus aureus : Proof of Concept. ACS Omega 7:46213–46221.

32. Anonymous. 2022. Clinical and Laboratory Standards Institute, M100: Performance Standards for Antimicrobial Susceptibility Testing. Table 2C.

33. Pettersen EF, Goddard TD, Huang CC, Couch GS, Greenblatt DM, Meng EC, Ferrin TE. 2004. UCSF Chimera--a visualization system for exploratory research and analysis. Journal of Computational Chemistry 25:1605–12.

34. Korb O, Stützle T, Exner TE. PLANTS: Application of Ant Colony Optimization to Structure-Based Drug Design, p 247–258. In (ed), Springer Berlin Heidelberg,

35. Korb O, Stutzle T, Exner TE. 2009. Empirical scoring functions for advanced protein-ligand docking with PLANTS. J Chem Inf Model 49:84–96.

